# Enrichment of spermatogonial stem cells and staging of the testis cycle in a dasyurid marsupial, the fat-tailed dunnart

**DOI:** 10.1101/2024.10.19.616824

**Authors:** Gerard A. Tarulli, Patrick R.S. Tatt, Rhys Howlett, Sara Ord, Beth Shapiro, Stephen R. Frankenberg, Andrew J. Pask

## Abstract

There is increasing interest in use of marsupial models in research, for use in next-generation conservation by improving fitness through genetic modification, and in de-extinction efforts. Specifically this includes dasyurid marsupials such as the Thylacine, Tasmanian devil, quolls and the small rodent-like dunnarts. Technologies for generating genetically modified Australian marsupials remains to be established. Given the need to advance research in this space, the fat-tailed dunnart (*Sminthopsis crassicaudata*) is being established as a model for marsupial spermatogonial stem cell isolation, modification and testicular transplantation. This species is small (60-90mm body size), polyovulatory (8-12 pups per birth), and can breed in standard rodent facilities when housed in a 12:12 light cycle. To develop the fat tailed dunnart as a model for next-generation marsupial conservation, this study aimed to enrich dunnart spermatogonial stem cells from whole testis digestions using a fluorescent dye technology and fluorescence-activated cell sorting. This approach is not dependent on antibodies or genetic reporter animals that are limiting factors when performing cell sorting on species separated from human and mouse by large evolutionary timescales. This study also assessed development of spermatogonia and spermatogenesis in the fat-tailed dunnart, by making the first definition of the cycle of the seminiferous epithelium in any dasyurid. Overall, this is the first detailed study to assess the cycle of dasyurid spermatogenesis and provides a valuable method to enrich marsupial spermatogonial stem cells for cellular, functional and molecular analysis.

## Introduction

The fat-tailed dunnart (*Sminthopsis crassicaudata)*, a mouse-sized marsupial from south-eastern Australia, provides several benefits over existing models in the study of spermatogonial stem cells. Other than its small size that makes for convenient housing and husbandry, spermatogenesis in the dunnart has several characteristics making it unique. Firstly, the anatomy of the testis is relatively simple, containing only 1-2 tubules with simple connections to collecting structures [1]. Secondly, dunnart testis tubules have the largest diameter of any known species, even amongst other dasyurids [1], and are 2-3 times wider than in the laboratory mouse though half as long. This makes handling tubules in experiments more straightforward. These features make the dunnart an attractive model to investigate the regulation of spermatogonial stem cells (SSCs) in unique ways. To enable the use of the fat-tailed dunnart as a model in this regard, we have established a technique to allow for enrichment of dunnart spermatogonia for use in downstream culture and molecular analysis. Furthermore, we have defined the stages of spermatogonial development, and characterised the cycle of the seminiferous epithelium in the fat-tailed dunnart, to identify features shared and distinct with/from eutherians.

Enriching spermatogonia from single cell suspensions broadly relies on the use of transgenic reporter models and/or antibodies specific for cell surface markers that are amenable to fluorescence-activated cell sorting (FACS). Extracellular proteins exhibit relatively rapid deviation during evolution, making the application of antibody-based FACS sorting difficult in species other than human, mouse and a few closely related species. Marsupials are separated from human and mouse by roughly 180 million years of evolution. Therefore, currently available antibodies raised to cell surface proteins relevant to germ cell research are not widely applicable for FACS-based enrichment of spermatogonia in marsupials. An alternative approach is to use cell-permeable dyes that fluoresce in response to the presence of specific proteins within specific cell types of the testis. One such molecule is BODIPY-aminoacetaldehyde, that fluoresces when acting as a substrate for aldehyde dehydrogenase (ALDH) proteins. ALDH enzymes are responsible for the conversion of retinal to retinoic acids that have 20-fold greater biological action and are critical in multiple steps during spermatogenesis [2]. Undifferentiated spermatogonia are dependent on ALDH expression by Sertoli and meiotic germ cells to switch from self-renewal to differentiation [3] and enter cycles of the seminiferous epithelium. The expression of ALDH enzymes is cell-type specific in the seminiferous epithelium and is known to be high in Sertoli and Leydig cells, as well as meiotic and post-meiotic germ cells, but low in early spermatogonia [4]. Thus, the use of ALDEFLUOR should enrich dunnart SSCs from single cell suspensions without the need for antibodies specific for these SSCs.

Of the 19 known human ALDH isoforms, 9 have been found to be detectable by ALDEFLUOR [5]. In mouse testis, four isoforms Aldh1a1, Aldh1a2, Aldh1a3 and Aldh8a1 are known to be expressed, and of these Aldh1a1, 1a2, and 1a3 can strongly induce ALDEFLUOR fluorescence. [4, 5]. ALDH expression in the mouse testis varies by cell type. Leydig cells strongly express all four isoforms, Sertoli cells express Aldh1a1 and Aldh8a1, spermatogonia only express Aldh8a1, preleptotene, leptotene and zygotene spermatocytes express Aldh1a3, pachytene and diplotene spermatocytes and meiotic cells express Aldh1a2 and Aldh1a3, round spermatids express only Aldh1a2 and elongating spermatids express Aldh1a1, Aldh1a2 and Aldh8a1 [4]. Based on these works, we hypothesise that in dunnart testis ALDH activity will be lowest in spermatogonia, as in mouse these cells only express Aldh8a1 that can only weakly activate ALDEFLUOR fluorescence.

Staging of the seminiferous epithelial cycle has been determined in many eutherians including the rat (*Rattus norvegicus* - 14 stages [6], mouse (*Mus musculus* - 12 stages [7], bovine (*Bos taurus* - 12 stages [8]), dog (*Canis familiaris* - 8 stages [9, 10]) and horse (*Equus caballus* - 8 stages [11]). The seminiferous epithelial cycle has also been determined in several marsupials including: the brush-tailed possum (*Trichosurus vulpecula*) [12, 13]), South American White-belly Opossum *Didelphis albiventris*), and the southeastern four-eyed opossum (*Philander frenatus*) [14] all with 10 stages; The Koala and southern hairy-nosed wombat (*Lasiorhinus latifrons*) [15], common ringtail possum (*Pseudocheirus pergrinus*)[16], musky rat-kangaroo (*Hypsiprymnodon moschatus*)[17], tammar wallaby (*Macropus eugenii)*, red-necked wallaby (*Macropus rufogriseus*), and long-nosed bandicoot (*Perameles nasuta*)[18] all observed to have 8 stages. It has previously been shown showed that while fundamentally similar in cellular associations, spermatids of some marsupials undergo morphological changes unique from eutherians. These changes include: flattening of the nucleus at right angles relative to the length of the elongating flagellum, the rotation of the nucleus at the point of connection to the flagellum and the incomplete coverage of the nucleus by the acrosomal cap [13, 15, 19]. These unique marsupial features, as well as dasyurid-specific characteristics described above, suggest that unique mechanisms of germ cell regulation can be revealed through a more detailed interrogation of spermatogenesis in the fat-tailed dunnart.

## Materials & Methods

### Collection and Processing of dunnart adult testis

Fat-tailed dunnart samples were obtained from a breeding colony maintained at the University of Melbourne, School of BioSciences. All animal handling and sample collection were approved by The University of Melbourne Animal Ethics Committee (#10206). Vitamin-supplement Pentavite was added to water at a ratio of 1:100 Pentavite:water. Animals were fed a daily wet mixture of 51% beef mince cat food, 36% beef and lamb flavoured cat biscuits, 12.7% Wombaroo small carnivore food and 0.3% calcium carbonate. Animals were kept between 21-25°C under a 16:8-hour light:dark cycle. Adult dunnarts killed for sample collection were euthanised by CO2 inhalation before cervical dislocation. Testis samples were punctured with fine forceps before fixation in Bouin’s solution (Merck). Samples were fixed overnight at 4°C then washed at room temperature rotating for 30 minutes in Phosphate-buffered saline (PBS) twice before storage in 70% ethanol at 4°C before processing for paraffin embedding. Sections for hematoxylin & eosin analysis were processed as described previously [20].

### Testis digestion & FACS for ALDH activity

Testes were dissected in 1:1 Dulbecco modified Eagle medium and Ham F12 (DMEM-F12), removing the tunica and placing the decapsulated seminiferous tubule in digestion mix containing collagenase type 1A (0.5 mg/ml) and DNase1 (0.65 mg/ml) in DMEM-F12 with 10% fetal bovine serum. Digestion occurred at 37°C for 15 minutes inverting every 5 minutes and ceasing whena single cell suspension was achieved. Samples were then passed through a 40 μm mesh filter before centrifugation at 4°C for 5 minutes at 850 rpm All DMEM-F12 medium contained 20mM HEPES buffer. The digested single cell suspension was diluted to one million cells/mL in ALDEFLUOR™ assay buffer (Stem Cell Technologies) and divided between tubes, including a control containing DEAB inhibitor, as a negative control for the ALDEFLUOR™ reagent and to determine gates for FACS. Tubes were incubated with 2 μL/mL of ALDEFLUOR™ reagent at 37°C for 45 minutes. The tubes were centrifuged at 1000 rpm, 4°C for 5 minutes before being resuspended in ALDEFLUOR™ assay buffer containing 0.2 μM DAPI (Sigma-Aldrich). Cells were sorted using a FACS-ARIA II (BD Biosciences), into 1.5 mL tubes containing DMEM-F12 medium supplemented with 10% FBS and 25 mM HEPES (Gibco) before cells were cultured *in vitro* or lysed using lysis buffer from the GeneElute Total RNA kit (Sigma Aldrich).

### In vitro culture of testis cell fractions and quantification of colony forming ability

Sorted and unsorted fractions were resuspended in DMEM-F12 medium supplemented with 10% FBS (Gibco), and 1% antimycotic/antibiotic (Gibco). Culture medium was changed every 3-4 days. For colony quantification, entire wells of a 12-well plate were imaged, and the total number of colonies counted. Examples of count images are shown in Supplementary Figure 1. Colonies composed of three or more round cells adhered to the primary feeder layer were used to determine colonies per cm^2^ in each of the sorted fractions after 20 days of culture. Data is presented as mean ±SD. Differences in colony-forming ability was analysed using two-tailed t-tests between all sorted populations.

### RNA Purification, Reverse Transcription & qPCR

RNA purification of sorted fractions was performed using the GeneElute Total RNA kit (Sigma-Aldrich) as per manufacturer’s instructions. Reverse transcription was performed using Superscript IV reverse transcriptase (Thermo Fisher Scientific) as per manufacturer’s instructions. Cell number input was standardised between samples for downstream qPCR analysis. qPCR was performed as described previously [21] using the PowerTrack Sybr reagent (Thermo Fisher Scientific) in a total of 10 uL. Gene-specific primers were based on a newly assembled dunnart genome [22] and designed to amplify regions spanning introns. Primers demonstrated a single curve by melt analysis. A control reaction without reverse transcriptase was used to verify primers were specific for cDNA. Primer sequences can be found in Supp. Table 1. Fold change was analysed using two-tailed t-tests between all sorted populations. Significance was set at a p-value of <0.05 and data expressed as mean ±SEM.

**Table 1.** Seminiferous tubule diameter of select marsupial and eutherian species (asterisks denote dasyurids)

| Species | Common Name | Tubule Diameter ($\mu\text{m}$) | Reference |
| --- | --- | --- | --- |
| **Present Study** |  |  |  |
| *Sminthopsis Crassicaudata* * | Fat-tailed dunnart | 452 ± 7 | This study |
|  |  | 510 | [1] |
| **Marsupials** |  |  |  |
| *Antechinus bellus* * | Fawn antechinus | 371.8 | [29] |
| *Antechinus bilarni* * | Sandstone false *antechinus* | 269.4 | [29] |
| *Antechinomys laniger spenceri* * | Kultarr | 460-520 | [1] |
| *Antechinus stuartii* * | Brown antechinus/Stuart's antechinus/ Macleay's marsupial mouse | 360-570 ± 75<br>460-520 | [1, 30] |
| *Dasyuroides byrnei* * | Kowari | 410-450 | [1] |
| *Didelphis virginiana* | Virginia opossum | 230-250<br>303 ± 7 | [11, 31] |
| *Lasiorhinus latifrons* | Southern Hairy-nosed Wombat | 244 ± 4 | [15] |
| *Macropus giganteus* and *Macropus fuliginosus* | Grey kangaroos | 180-320 | [32] |
| *Megaleia rufa* | Red kangaroo | 200-250 | [33] |
| *Phascolarctos cinereus* | Koala | 228 ± 6 | [15] |
| *Philander frenatus* | South-eastern four-eyed opossum | 279 ± 17 | [34] |
| *Sarcophilus harisii* * | Tasmanian devil | 360 | [1] |
| *Sminthopsis leucopus* * | White-footed dunnart | 430 | [1] |
| *Trichosurus vulpecula* | Common brushtail possum | 226-288 | [35] |
| **Eutherians** |  |  |  |
| *Bos grunniens* | Datong yak | 246 ± 1 | [36] |
| *Bos taurus* | Zebu bull | 257.0 ± 7 | [37] |
| | Holstein-Friesian | 228.04 ± 89 | |
| | HK | 183.43 ± 5 | |
| *Calotes versicolor* | Oriental garden lizard | 248.02 ± 3 | [38] |
| *Camelus dromedarius* | One humped camel | 202.3 ± 9 | [39] |
| *Capricornis crispus* | Tokara (Japanese native Goat) | 198 ± 1.0 | [40] |
| *Canis familiaris* | Dog | 143-245 ± 3 | [41] |
| *Cerdocyon thous* | Crab-eating fox | 236 ± 22 | [42] |
| *Cnemidophorus gularis* | Common spotted whiptail lizard | 260 | [43] |
| *Cophosaurus texanus* | Greater earless lizard | 355 | [43] |
| *Diphylla ecaudata* | Hematophagous bat | 195.1 ± 6 | [44] |
| *Equus asinus* | Donkey | 222 ± 6 | [45] |
| *Equus asinus* × *Equus caballus* | Mule | 127 ± 4 | [45] |
| *Felis catus* | House cat | 223 ± 5 | [46] |
| *Heterocephalus glaber* | Naked mole rat | 192 ± 21 | [47] |
| *Homo sapiens* | Human | 163.0 ± 9 | [48] |
|  |  | 227.9 ± 23 | [49] |
| *Leopardus tigrinus* | Oncilla (little spotted cat) | 228.29 ± 22 | [50] |
| *Mus musculus* | Mouse<br>C57Bl/6 | 216 ± 2 | [51] |
|  | Swiss | 228 ± 2 |  |
|  | Balb/c | 210 ± 3 |  |
| *Rattus norvegicus* | Albino rat | 265 | [52] |
|  |  | 319.8 ± 12 | [53] |
|  |  | 330 | [54] |
| *Rhea americana* | Greater rheas | 122.5 ± 30 | [55] |
| *Sceloporus aeneus* | Mexican endemic oviparous lizard | 319.1 | [56] |
| *Sus scrofa* | Boar | 280 ± 20 | [57] |
| *Ursus arctos horribilis* | Grizzly bear | 165.23 ± 23.6 | [58] |

## Results

### Testicular tubule diameter

The most notable feature of the dunnart testis, and of several dasyurid species including a large number belonging to the *Antechinus* genus, is having the largest tubule diameter of any known closely linked group of species [1]. Replicating early studies in the dunnart and compiling more recently published data in a broad range of species (Table 1), no eutherian species with a larger diameter has been found, to the date of this publication. This is not a feature common in marsupials more generally, as other Australian marsupials possess diameters much closer to those of humans and mice.

### Cycle of the seminiferous epithelium

The seminiferous epithelial cycle in the fat-tailed dunnart can be categorised into 12 stages based upon changes in round spermatid acrosome and nuclear morphology (stages I-X), as well as stages of spermatocyte development (XI-XII) (Figure 1). A conceptual summary of the cellular associations in each stage of dunnart spermatogenesis can be found in Figure 2

**Figure 1.**
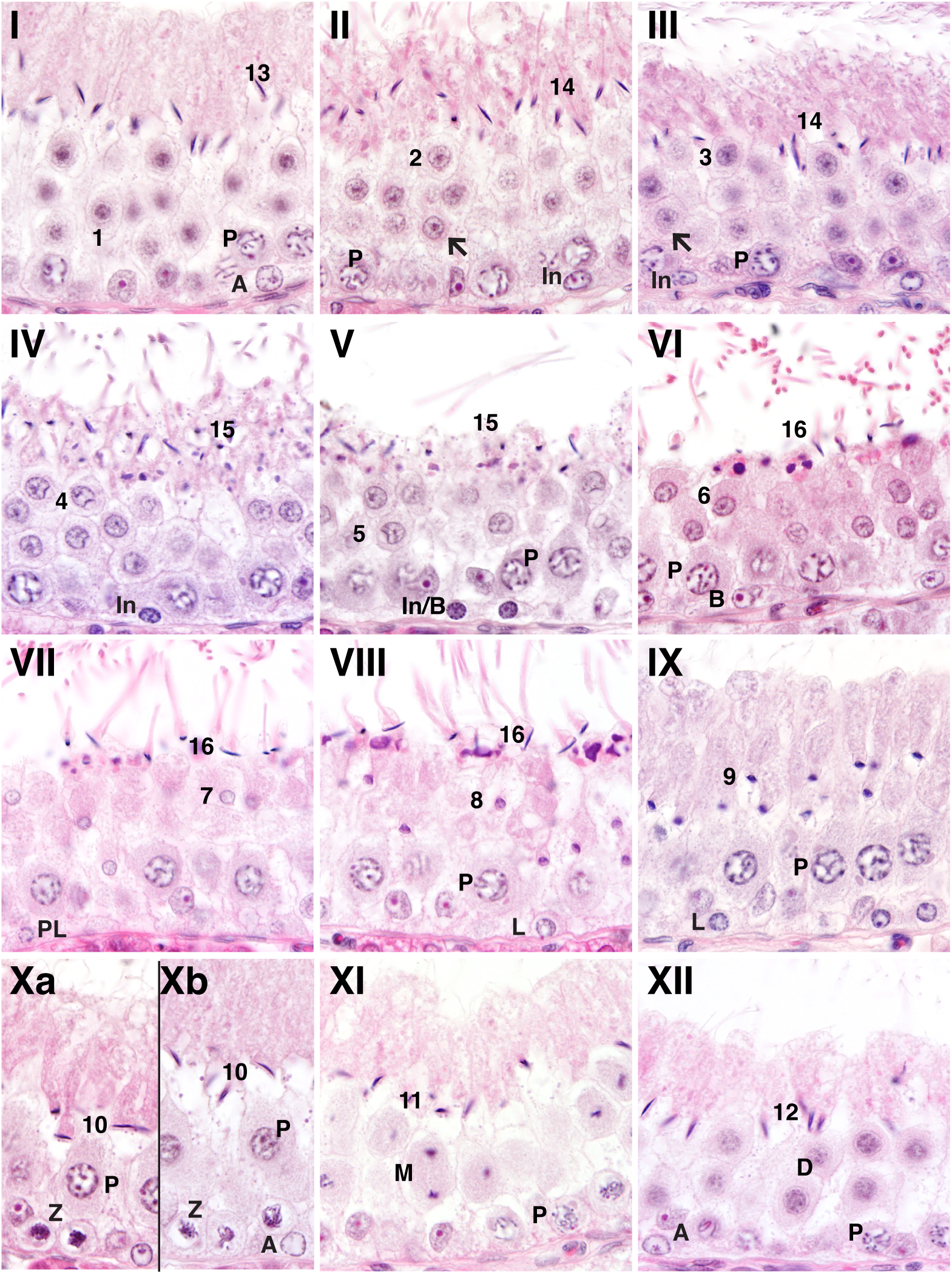
Twelve stages of the seminiferous epithelial cycle during dunnart spermatogenesis. The dunnart has a protracted process of spermiation, occurring over 4 stages coincident with significant round spermatid nuclear compaction. Stages are denoted with roman numerals. See text for detailed description of changes in cellular morphologies. (A = A spermatogonium, In = Intermediate spermatogonia, B = B spermatogonia, PL = preleptotene spermatocyte, L = Leptotene spermatocyte, Z = Zygotene spermatocyte, P = pachytene spermatocyte, D = Diplotene spermatocytes, M = mitotic spermatocyte, R = Round spermatid, 1-16 = spermatid steps).

**Figure 2.**
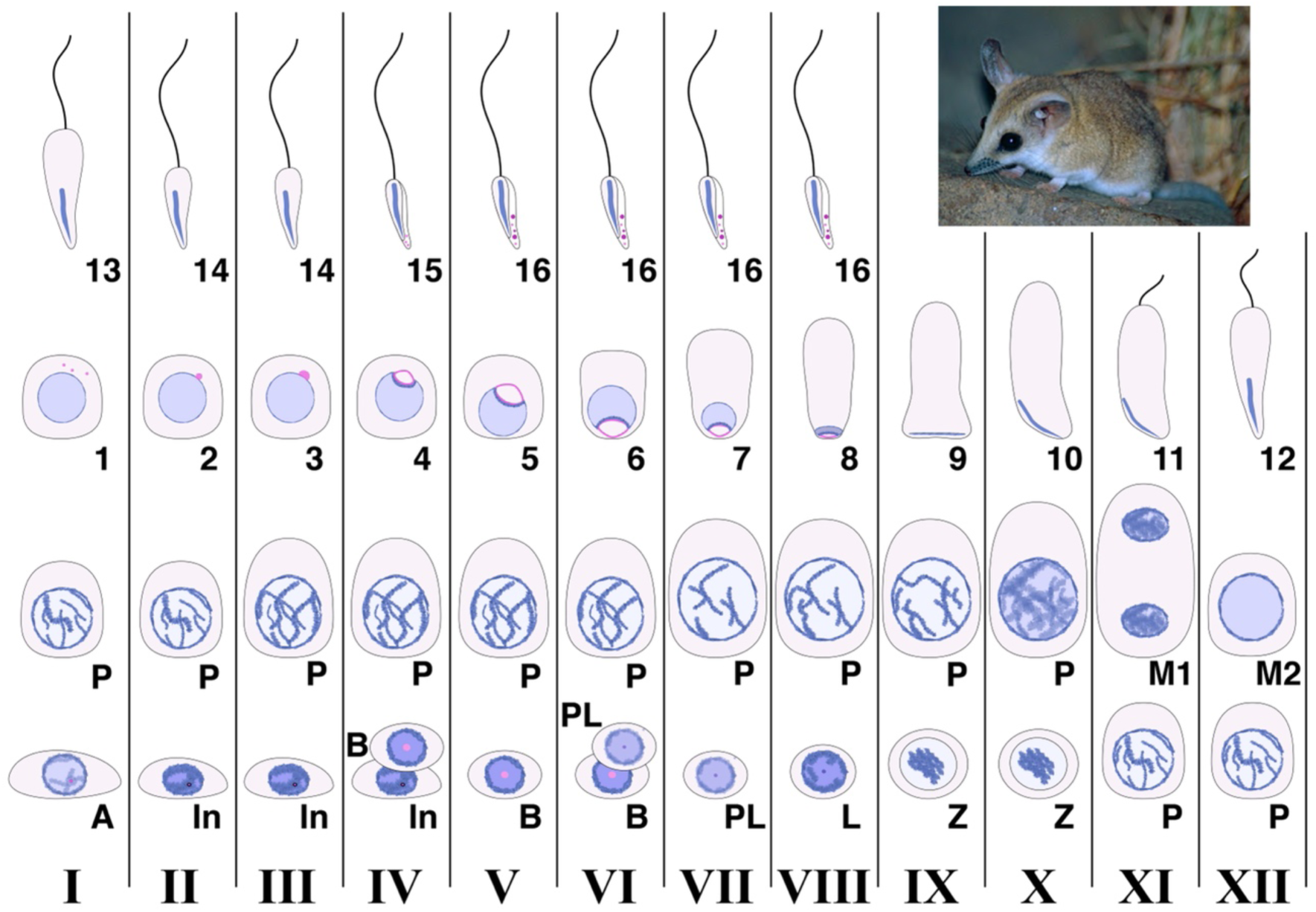
Conceptual diagram of cellular associations found in the 12 stages of the cycle of dunnart seminiferous epithelium. A = A spermatogonium, In = Intermediate spermatogonia, B = B spermatogonia, PL = preleptotene spermatocyte, L = Leptotene spermatocyte, Z = Zygotene spermatocyte, P = pachytene spermatocyte, D = Diplotene spermatocytes, M = mitotic spermatocyte, R = Round spermatid, 1-16 = spermatid steps).

### Spermatogonia

*A* spermatogonia are found in all stages along the basement membrane, and clearly distinguishable from other cells occupying this niche, particularly at stage I. *A* spermatogonia possess a large spherical to semi-ovoid nucleus, with thin nuclear strands apparent around nuclear edges. A second population of spermatogonia distinguished by a smaller ovoid cytoplasm, ovoid-to-round nucleus and moderate to darkly-stained chromatin, are visible through stages II-IV and classified as Intermediate (*In*) spermatogonia. An example in Figure 1, Stage V shows two *In/B* spermatogonia adjacent to the basement membrane. Nuclear and cytoplasmic morphology of these cells is distinct from *A* or *B* spermatogonia that have a pale nuclear appearance (see Stage V). *B* spermatogonia have spherical nuclear morphology with a prominent nucleolus and chromatin staining paler than found in *In* spermatogonia.

### Stage I

Step 1 round spermatids from the second meiotic division are newly formed at this stage, with no evidence of an acrosome. Condensed chromatin strands are observed in pachytene spermatocytes that are also apparent at many other stages of the cycle. The Sertoli cell nucleus has a curved trapezoid morphology, more closely resembling the human rather than rodent nuclear morphology.

Step 13 elongated spermatids from the previous cycle of spermatid development populate the most luminal layer of the seminiferous epithelium, with the heads oriented perpendicular to the basement membrane. Significant spermatid cytoplasm is still apparent extending near to the lumen, with a relatively short tail evident. Pachytene spermatocytes have clearly defined chiasmata and are increasing in size compared with previous stages of development. *A* spermatogonia are present at this stage, with characteristic large round nucleus with pale staining chromatin.

### Stage II

Step 2 spermatids are defined by the presence of a small acrosome that has just made contact with the nuclear membrane, without lateral spreading and condensation of nuclear architecture. Nuclear position remains central in the cell. Step 14 spermatids have an elongated flagella extending into the tubule lumen, with a reduced proportion of cytoplasm in basal aspects of these cells. Pachytene spermatocytes continue to increase in size. *In* spermatogonia, defined by an ovoid nucleus with a rim of condensed chromatin as well as ovoid cytoplasm, are frequently observed at this stage and reside close to the basement membrane. Intermediate spermatogonia are evident from this stage.

### Stage III

Step 3 spermatid acrosomes have contacted the nuclear membrane and grown in size, and spreading across the membrane is evident (arrow). *In* spermatogonia are still present at this stage. Pachytene spermatocytes increase in size compared with Stage II.

### Stage IV

Step 4 spermatids have a clearly defined nuclear indentation at the site of acrosomal fusion. Though the nucleus is still centrally located within the cell. Step 15 spermatids have a further reduced cytoplasm and contain eosin-staining inclusions within the basal cytoplasm. There is further elongation of flagella. *In* spermatogonia are present along the basement membrane, defined by an ovoid cell and nuclear morphology.

### Stage V

The nucleus of Step 5 round spermatids is still centrally located within the cytoplasm, and the acrosome has spread across the nucleus to indent a width close to that of the nucleus, though still only occupying 20-30% of the nuclear area. Removal of cytoplasm in Step 15 spermatids is almost complete at this stage. Spermatogonia with characteristics of both *In* and *B* subtypes are present along the basement membrane.

### Stage VI

Stage VI is the first stage to exhibit residual body formation and apparent initiation of spermiation in Step 16 spermatids. The presence of similar elongated spermatid morphology until Stage VIII, demonstrates a protracted process of spermiation in the fat-tailed dunnart.

The Step 6 spermatid nucleus begins to become basally oriented in the cell, though no reductions in nuclear size is apparent at this stage. The acrosome of the nucleus is not yet oriented toward the basement membrane. *B* spermatogonia are present along the basement membrane, characterised by a round nucleus with large central nucleolus and dense chromatin at the nuclear edges.

### Stage VII

Step 7 spermatids show a reduction in nuclear volume and orientation of the acrosome to the basement membrane. Preleptotene spermatocytes are still observed along the basement membrane. Preleptotene spermatocytes continue to be found at this stage. Step VII is where the first of pre-leptotene spermatocytes appear, which are discernible by their relatively small cellular and nuclear size, and diffuse nuclear staining.

### Stage VIII

Step 16 spermatids undergo spermiation, leaving large, darkly stained residual bodies on the luminal edge of Sertoli cells. Compaction of chromatin is first apparent in Step 8 spermatids, with the luminal side of spermatids more condensed than the basal side. Leptotene spermatocytes are observed on the basement membrane, defined by a nuclear and cellular size greater than preleptotene spermatocytes, and sharing a similar diffuse pattern of chromatin.

### Stage IX

The Step 9 spermatid nucleus is further compacted, and chromatin condensed. The cytoplasm of these cells extends outwards from the basally-oriented nucleus as the cytoplasmic length increases. Pachytene spermatocytes size is the greatest of any stage. Leptotene spermatocytes are still present along the basement membrane.

### Stage X

Step 10 spermatid nucleus becomes oriented parallel with the basement membrane. This is a unique feature found in a limited number of other marsupials, particularly dasyurids. Zygotene spermatocytes are first observed at Stage X, with small, highly condensed chromatin strands though still occupying a basal position within the epithelium. At a later point of Stage X (Figure 1, Xb), the spermatid nucleus begins to re-orient to be perpendicular to the basement membrane. Zygotene spermatocytes are present and identified by condensed chromatin have moved from the basement membrane. The chromatin of pachytene spermatocytes begins to become more diffuse at this stage, with an increase in cell size.

### Stage XI

Step 11 spermatid cytoplasm reduces in length, along with the size of the spermatid head. This is found in association with the earliest evidence of a flagellum. Pachytene spermatocytes undergo the first meiotic division at this stage, and the nuclear size and re-orientation of chromatin in zygotene spermatocytes from Stage X increases to mark the initiation of differentiation into early pachytene spermatocytes (Figure 1, XI, Z/P).

### Stage XII

Step 12 spermatid cytoplasmic length continues to decrease, with clear flagella that span less than 50% of total cell length. Secondary spermatocytes are at the diplotene stage, with diffuse chromatin filling a spherical nucleus. These cells undergo the second meiotic division to become Step 1 round spermatids at Stage I.

### Fluorescence-activated cell sorting to enrich for dunnart testicular cell types

We invested significant time finding antibodies that could be used for live cell FACS of dunnart testicular cell suspensions, however this was unsuccessful. This is likely due, at least in part, to the evolutionary distance between marsupials and human/mouse, and the relatively fast pace of evolution in genes encoding extracellular proteins. An alternative approach is using live cell-permeable fluorescent dyes. One such dye is ALDEFLUOR, a BODIPY-aminoacetaldehyde that acts as a substrate for aldehyde dehydrogenase (ALDH) proteins that are expressed only at low levels in spermatogonia but higher in meiotic and post-meiotic germ cells, as well as in major somatic cell types of the testis. To verify the likelihood that ALDEFLUOR will show the same cell type-specific expression in dunnart as in mouse testis, we assessed the genetic phylogeny of major ALDH genes (i.e. those shown to activate ALDEFLUOR fluorescence and that are expressed in mouse testis [4, 5]). Predicted ALDH coding sequences were identified by BLAST searches in a newly assembled dunnart genome (Frankenberg et al., unpublished) and aligned with mouse ALDH genes. These ALDH genes (*ALDH1A1*, *1A2* and *1A3*) belong to the same clade in mouse and dunnart (Figure 3A), aside from an additional ALDH gene in mouse (*Aldh1a7*) that segregates with mouse *Aldh1a1* and forms a separate clade to two dunnart *ALDH1A1*-like paralogues, which we refer to as *ALDH1A1-A* and *ALDH1A1-B*.

**Figure 3.**
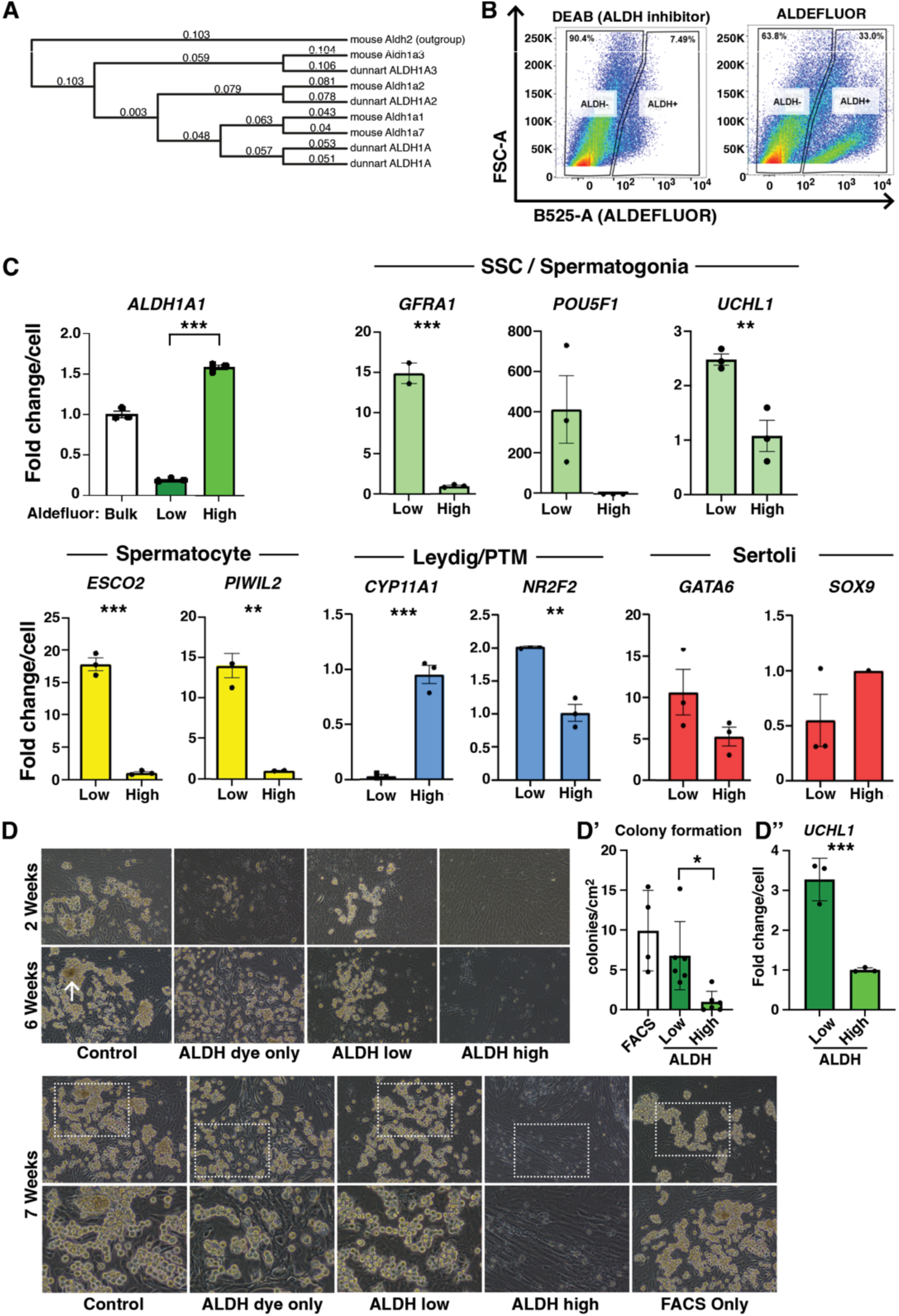
Enrichment and survival of dunnart spermatogonial stem cells using ALDEFLUOR. A) Genetic phylogeny of dunnart and mouse ALDH genes. B) FACS profiles of dunnart testis single cell suspensions treated with control (ALDEFLUOR plus DEAB, left) and ALDEFLUOR only (right). C) qPCR analysis of cell subtype-specific genes in sorted ALDH-low and -high cells. D) Sorted ALDH-low cells survived and proliferated into grape-like clusters after 6 weeks in culture, while little germ cell growth was observed in the ALDH-high sorted fraction. D’) Quantification of colony formation. D’’) qPCR analysis of UCHL1 expression in ALDEFLUOR-sorted cells, cultured for 7 weeks. Asterisks denote levels of significance (* <0.05, ** <0.01, ***<0.001).

Nonetheless, the *ALDH1* genes between mouse and dunnart show close evolutionary associations compared with other ALDH genes, highlighting the likelihood that ALDEFLUOR fluorescence reflects ALDH activity in the same way in dunnart. In testing ALDEFLUOR on dunnart cells, testes from adult dunnarts were digested to single cell suspensions, labelled with ALDEFLUOR and sorted into two populations based on levels of fluorescence compared with the ALDH inhibitor, DEAB (Figure 3B). Cells were sorted for RNA extraction and qPCR analysis. Primers specific *ALDH1A1*, and for genes expressed by dunnart testicular somatic cell subtypes (Leydig - *CYP11A1*, Leydig/peritubular myoid cells - *NR2F2*, Sertoli cells – *SOX9*, *GATA6*) and germ cells (SSCs - *POU5F1*, SSCs/spermatogonia - *GFRA*; differentiating germ cells - *ESCO2*, *PIWIL2*) were developed and tested on ALDH-high and - low fractions (Figure 3C). As expected, ALDH-high cells expressed greater *ALDH1A1* levels compared with ALDH-low cells. Significant enrichment of markers specific for SSCs, spermatogonia and spermatocyte was observed in ALDH-low cells. ALDH-low cells expressed very little of the Leydig cell marker *CYP11A1*, demonstrating that ALDEFLUOR is able to enrich for spermatogonia while excluding Leydig cells with high efficiency. However, Sertoli cell markers *GATA6* and *SOX9* were not enriched in ALDH-low or -high cells, and a marker of peritubular myoid cells, *NR2F2*, was enriched 2-fold in ALDH-low cells. Altogether this demonstrates that while ALDEFLUOR can successfully enrich for early germ cell types - including SSCs - and exclude Leydig cells, it is not able to efficiently exclude Sertoli cells or peritubular myoid cells.

To determine whether digestion and sorting by ALDH can enrich for cells with the potential to form colonies in vitro, unsorted and ALDH-sorted cells were cultured for up to 7 weeks (Figure 3D). Within 2 weeks small colonies of germ cells formed in ALDH-low sorted samples, as well as control and dye-only samples. At 18 days, colony numbers were counted (Figure 3D’), finding a x.x-fold increase in colony forming ability in ALDH-low sorted samples. Within 6 weeks clear expansion of these colonies is observed (note images are of the same region at each timepoint in Figure 3D). The colonies in control wells included large cellular aggregates and structures resembling testicular cords (Figure 3D, arrow in 6-week control). In stark contrast very few colonies established in ALDH-high samples by this time. Further, when compared to cells sorted in the absence of ALDEFLUOR, that better reflects stresses of handling, fewer cellular aggregate established in ALDH-low sorted cells. Rather, cells grow more often as chains connecting smaller clusters than in FACS-only cells. Of note, ALDH-high sorted cells formed a confluent sheet of cells by 4 weeks, that sloughed from the plastic surface, and then reseeded additional layers of cells. This phenomenon was not observed in ALDH-low sorted cells. Assessment of SSC marker expressed showed a 3-fold increase UCHL1 in ALDH-low cultured samples (Figure 3D’’)

In an attempt to overcome contamination with Sertoli and peritubular somatic cells, a more refined gating strategy was tested that differentiated FACS plots into 5 sub-populations (denoted in red numbers in Figure 4A) based on ALDEFLUOR fluorescence and an amplified forward-scatter channel, where five gated populations were discernible (denoted in red numbers in Figure 4A). These were used to enrich for spermatogonia expected to be present in gates 1 and/or 2 where ALDH activity is low. Gates 3-5 are expected to enrich for other cell types. Cells were sorted and assessed for cell-specific markers by qPCR (Figure 4B). Enrichment for *ALDH1A1*-expressing cells by ALDEFLUOR was consistent with previous experiments. The Leydig cell-specific marker, *LHR*, was enriched in populations 4 and 5 that shows intermediate or high ALDH activity, respectively, while the Sertoli cell-specific markers *SOX9* and *GATA6* were enriched in populations 2, 4 and 5 to similar extents. Populations 1 is low for all markers assessed and likely consist of haploid spermatids and other interstitial cell types including immune cells that likely also contribute to population 3. Overall, a more refined gating approach did not further exclude Sertoli cells from the ALDH-low sorted population

**Figure 4.**
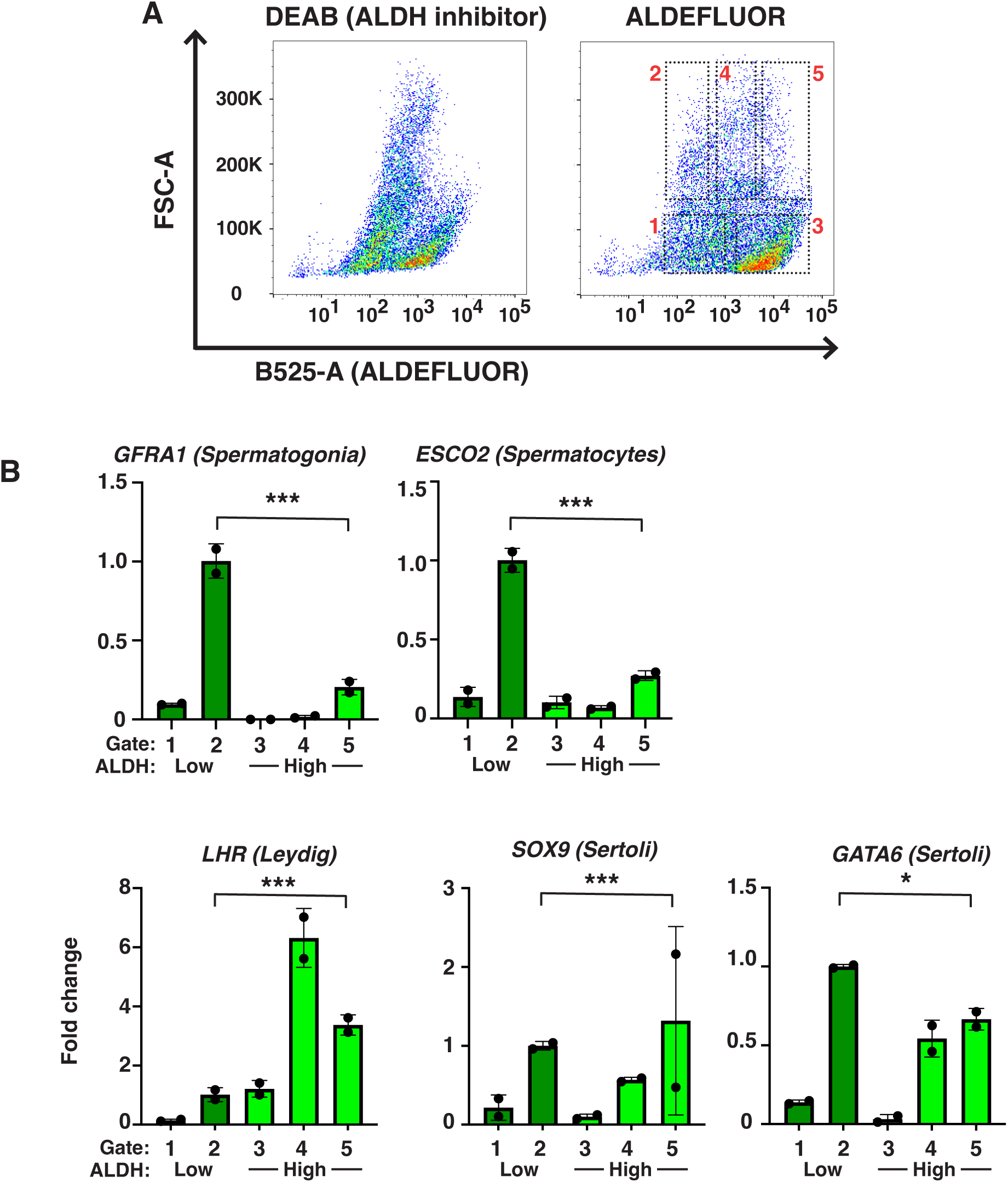
Refined gating of ALDH subpopulations does not improve exclusion of somatic cell types. A) An improved gating strategy allows for distinction of five populations based on ALDEFLUOR fluorescence and forward scatter. Populations 1-2 are ALDH-low and 3-5 are ALDH-high. B) qPCR analysis of sorted populations demonstrates a significant enrichment of spermatogonia and spermatocytes in ALDH-low population 2. Leydig cells are enriched in ALDH-high populations 4 and 5, while Sertoli cells are enriched in populations 2, 4 and 5. Asterisks denote levels of significance (* <0.05, ** <0.01, ***<0.001).

## Discussion

This study is the first to provide a means to enrich for marsupial spermatogonia from digested testicular tissue, using a cell-permeable dye, ALDEFLUOR. Results show that ALDEFLUOR is able to enrich markers of dunnart spermatogonia up to 15-fold, and colony forming ability >30-fold. With the absence of contemporary technologies to enrich SSCs from marsupial testis digests, such as reporter models and SSC-specific antibodies, ALDEFLUOR is a valuable tool to move research forward in marsupial SSC biology. For example, that spermatogonial-enriched fractions survive and robustly expand in long-term culture provides the opportunity to genetically modify and transplant SSCs to generate marsupial reporter models. This furthers the broader goal of developing marsupial genetic rescue technologies.

The use of the ALDEFLOUR dye and FACS in this study allowed for enrichment of SSCs and early spermatocytes, as well as exclusion of Leydig cells from primary dunnart testis digests. However the SSC-enriched fraction still contained somatic cell populations, in particular Sertoli and peritubular myoid cells. Nonetheless, the very low colony-forming ability of ALDH-high sorted cells indicates that these cells may be isolated using differential plating/serial passaging. This first successful use of ALDEFLUOR in a marsupial species makes use in other species likely to be equally successful, given that marsupials and eutherians are segregated by roughly 180 million years of evolution. ALDEFLUOR represents a straightforward way to enrich spermatogonia/SSCs from any species where conventional approaches (antibody labelling and/or reporter models) are not available. While the technique is fast and straightforward, it suffers from relatively low accuracy of enrichment compared with mouse reporter models, particularly when used in combination with antibodies specific for spermatogonia. This could be due to the stage-specific expression of Sertoli cell ALDH, required for the orchestration of spermatogonial differentiation during the epithelial cycle, that results in a subpopulation of Sertoli cells having lower ALDEFLUOR fluorescence. This may be overcome with stricter gating strategies and the identification/development of antibodies able to specifically bind cell surface proteins on dunnart spermatogonia. On this latter point, improvements in the quality of a dunnart transcriptome will enable the development of marsupial-reactive antibodies to key cell surface markers on dunnart SSCs. Further, using ALDEFLUOR to partially enrich SSC populations will be useful in facilitating the identification of SSC-specific genes in dunnarts, through the assessment sorted cell transcriptomes at single cell resolution. These avenues provide fertile ground for future study. This is particularly true in light of previous attempts to use ALDEFLUOR with/without additional enrichment techniques to isolate mouse SSCs [24, 25]. In these studies, lack of ALDEFLUOR signal alone was unable to significantly enrich for transplantation-competent germs cells. However, when used in combination with either CD9- or CDH1-specific antibodies a lack of ALDEFLUOR greatly increased testis colony formation after transplantation [24].

The present study also collated data on seminiferous tubule diameters in different species, and showed that members of the dasyurid family possess the largest tubule diameter of any mammalian species assessed to date. This is an important finding for plans to use the fat-tailed dunnart as a surrogate for de-extinction of the Tasmanian tiger, and in ongoing efforts to develop functional approaches for genetic rescue of other dasyurid marsupials (e.g. northern quoll). This tubule diameter is likely to aid in success of SSC tubule injection, due to the relatively large target area within the dunnart testis.

This study is the first to define the stages of spermatogenesis in the dunnart, or any dasyurid, and identify a relatively protracted process of spermiation that may be unique to this species, or marsupials more generally. Results of this study identified 12 distinct stages of the dunnart seminiferous epithelial cycle, that is larger than the number identified in most other marsupials. It is possible that the smaller number of stages defined in previous studies of spermatogenesis in some marsupials (see below) may be partly explained by the difficulty in using spermiation to classify tubules as being in Stage VIII. In some marsupials, using this parameter alone to define Stage VIII tubules may identify more than one stage when additional defining parameters are applied. The cellular associations observed in the dunnart seminiferous epithelium are fundamentally akin to associations observed in eutherian species and other marsupials studied to date [12, 15–18, 26–28], even within sub-populations of spermatogonia. The major difference apparent in the development of spermatids at stage X where the spermatid nucleus is arranged perpendicular to the long axis of the cell, and the incomplete coverage of the nucleus by the acrosomal cap. No direct mechanisms have been found to explain the unique spermatid nuclear organisation found in marsupials, however it is unlikely to be associated with the large tubule diameter of the dunnart seminiferous epithelium, as this feature is also found in non-dasyurid marsupials that have tubular diameters closer to mouse and human [15, 17].

Overall this study provides a clear framework upon which future studies of dunnart spermatogenesis can be interpreted, along with a tool to enrich spermatogonia from digested primary testis tissue, without the need for antibodies specific for cell-surface markers. This knowledge can also be applied in efforts to target marsupial spermatogonial stem cells for genetic modification in the context of current conservation and de-extinction efforts.

## Acknowledgements

The authors wish to give credit to Dr. Vanta Jameson & Dr Magdaline Sakkas from The University of Melbourne cytometry core facility at the Melbourne Brain Centre for assistance with cell sorting, Bioresources staff maintaining dunnart husbandry, and members of the Biological Optimal Microscopy Platform for maintenance of microscopy hardware.

## Data availability

The data underlying this article will be shared on request to the corresponding author.

