## Supplementary figures and images for "Enrichment of spermatogonial stem cells and staging of the testis cycle in a dasyurid marsupial, the fat-tailed dunnart"

### Supplementary Figure 1

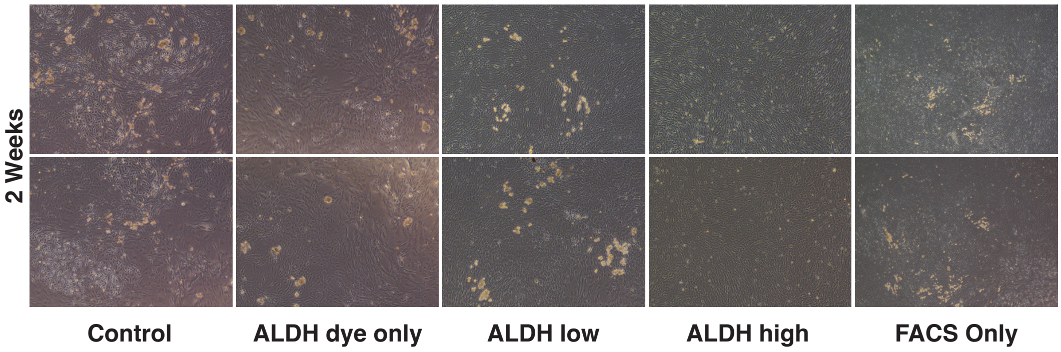


***Supp. Figure 1 – Example of images used to quantify colony forming ability in Figure 3D'***
