## Supplementary Table 1 for "Enrichment of spermatogonial stem cells and staging of the testis cycle in a dasyurid marsupial, the fat-tailed dunnart"

| Primer | Sequence |
|---|---|
| *SOX9* | F - CAATGGCTCTAGCAAGAACAAGC<br>R - CGTTTCTCACTCTCATTTAACAGCC |
| *POU5F1* | F - TCAGTAATATTGCTGAGGAGCTGG<br>R - CATAAGTGGGATGACTGAATGGAGG |
| *GATA6* | F - GAATTCAAACAAGGAAGCGAAAACC<br>R - TGTGTTGTAGGAGAAGCATTTTTGC |
| *UCHL1* | F -<br>R - |
| *GFRA1* | F - TAAAATCCAACGCTTCAGGAAATCC<br>R - TCTGCCAGACTTATGAACAAGAAGG |
| *PIWIL* | F - AAACCTGAGTCCTGACCATATGC<br>R - ATGATGTAAGACTTGTCCAGACAGG |
| *ESCO2* | F - TCTTTAACCTGATGAGAAGAAAGGGC<br>R - AAGTTGGGTGTATTGCAGTATTTGG |
| *CYP11A1* | F - ACTTCCGTGCTCTGAGCTTT<br>R - AAGGTGAACACGATGGGCTT |
| *NR2F2* | F - TGGAAAAACTGAAGGCATTACACG<br>R - GTACTGGCTCCTAACATACTCTTCC |
| *LHR* | F - AGTCACTGCTGTGCATTTATAAACC<br>R - TCCCAGATACTTAGTTCACTCTCCG |
| *ESCO2* | F - TCTTTAACCTGATGAGAAGAAAGGGC<br>R - AAGTTGGGTGTATTGCAGTATTTGG |
